# Learn to Track: Deep Learning for Tractography

**DOI:** 10.1101/146688

**Authors:** Philippe Poulin, Marc-Alexandre Côté, Jean-Christophe Houde, Laurent Petit, Peter F. Neher, Klaus H. Maier-Hein, Hugo Larochelle, Maxime Descoteaux

## Abstract

We show that deep learning techniques can be applied successfully to fiber tractography. Specifically, we use feed-forward and recurrent neural networks to learn the generation process of streamlines directly from diffusion-weighted imaging (DWI) data. Furthermore, we empirically study the behavior of the proposed models on a realistic white matter phantom with known ground truth. We show that their performance is competitive to that of commonly used techniques, even when the models are used on DWI data unseen at training time. We also show that our models are able to recover high spatial coverage of the ground truth white matter pathways while better controlling the number of false connections. In fact, our experiments suggest that exploiting past information within a streamline's trajectory during tracking helps predict the following direction.

## 1 Introduction

Tractography is currently at the heart of human brain connectomics studies [15]. However, recent biases and limitations of existing tractography pipelines have been highlighted [4], such as the reconstruction of many non-existent connections (false positive streamlines), poor spatial extent of existing connections and the difficulty of injecting anatomical priors beyond manual dissection and tissue classes from T1-weighted segmentations.

Currently, tracking algorithms depend on local models with assumptions on the nature of the underlying DWI signal. In 2015, [13] proposed a machine learning approach to fiber tractography based on a random-forest classifier. They successfully demonstrated how a purely data-driven approach can be used to reconstruct streamlines from the raw diffusion signal. Their method works well on 2D synthetic data and shows promising qualitative results on *in vivo* data. However, it has yet to be shown how well machine learning (and particularly deep learning) approaches can perform quantitatively on more realistic data and how well they can generalize to unseen data. In this paper, our main contributions are the first deep learning models for this problem and their evaluation, namely (1) a local reconstruction model based on a multilayer perceptron, (2) a sequential reconstruction model based on a recurrent neural network, (3) a careful quantitative evaluation of performances on the phantom of the ISMRM 2015 Tractography Challenge, and (4) a qualitative examination of the streamlines generated in unseen data during training. Our method outperforms or is competitive with the current state-of-the-art deterministic and probabilistic tractography algorithms robust to crossing fibers. In particular, out of 96 other tractography methods, this is the only approach able to recover more than 50% of spatial coverage of ground truth bundles while producing overreaching false connections below 50%. Our recurrent neural network is a promising deep learning solution for tractography based on raw DWI. It includes a notion of history of followed directions, which makes it robust to crossing fibers, robust to a wide range of geometries and allows the flexibility to include priors and learn how to reduce false-positive connections.

## 2 Using Deep Learning for Tractography

Given a diffusion dataset and sequences of spatial coordinates, the goal is to train a model to predict tracking directions to follow. In the context of tractography, a deterministic model can be used in an iterative process for streamline creation.

We chose to focus on deep learning models because of their well-known ability to discover and extract meaningful structures directly from raw data [9]. Our models are based on two types of deep learning models : a *Feed-Forward Neural Network* (FFNN), and a *Recurrent Neural Network* (RNN) [7]. While the FFNN is a local model and serves as a good baseline, it has the same weaknesses as existing methods, i.e. it is not able to learn streamline structures. To address this weakness, we used an RNN, because this family of models can process whole sequences as input. In our case, treating streamlines as sequences of coordinates in 3D space, our hypothesis is that a recurrent model should be able to learn the fiber or bundle structure through the diffusion signal in order to make better predictions and solve classic problems like fiber crossing.

*Model inputs:* As in [13], to be independent of the gradient scheme, the raw diffusion signal is first resampled to have *D* gradient encodings evenly distributed on the sphere (we used *D* = 100). We also normalized each diffusion-weighted images by the b=0 image. A streamline is represented as a sequence ***S*** of *M* equally-spaced spatial coordinates *P*_i_ = (*x_i;_y_i;_Z_i_*). The diffusion signal is evaluated at each of these points, using trilinear interpolation in the voxel space. This results in a sequence of *M* vectors with *D* dimensions representing the diffusion information along the streamline. In all our models, we also tried giving the previous direction as a supplementary input, as in [13]. Note that the spatial coordinates are not given as input to the model. This choice allows the model to be invariant to brain size or translation, reducing the preprocessing needed before feeding data to the model and improving generalization.

**Fig. 1.**
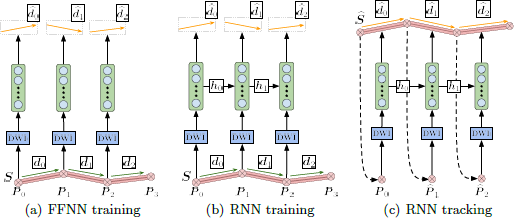
Architecture of the proposed models, (a) Given a streamline *S,* diffusion information is evaluated at each point *Pi* using trilinear interpolation **(DWI(Pi)).** The resulting vector is provided to the FFNN to predict a direction *di* (orange), which is compared against its associated target direction *di* (green), (b) Unlike the FFNN, the RNN has recurrence connexions through each step, allowing to send information to itself through the sequence, (c) Given a starting point **Po,** a new streamline *S* is generated by iteratively predicting a new direction *di,* and feeding the estimated new position *Pi+i* back to the model. Note how the predicted direction *di* gets influenced by prior information along the streamlines through *hj<i.*

### 2.1 Models

**FFNN** The FFNN sees all streamline coordinates as individual, independent local data points. The output of the model is a 3-dimensional normalized vector. The model is represented in Figure 1(a). To remove the directional ambiguity when no previous direction is given, we choose to consider the output vector as an *undirected axis* instead of a direction. To this end, the loss function is defined as the negative squared cosine similarity.

**RNN** The general idea behind the RNN is to model an internal state that is updated with each new observation in the input sequence and can be used to make predictions. Through its updatable internal state, the model can “remember” relevant features about the past. In this case, we used a Gated Recurrent Unit (GRU) [3] type of RNN.

Figure 1(b) shows that for each point *P_i_* in the streamline, the diffusion information **DWI**(*P_i_*) is used to update the internal state *h_i_* of the model. From there, at each step along the streamline, the model makes a prediction of the direction to follow 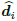. The loss function is defined as the mean squared error (MSE) between the model's prediction 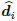 and the target *d_i_* (i.e. the next normalized segment of the streamline).

### 2.2 Tractography

Tractography is performed by using a fully trained model. Streamlines generation follows an iterative process as in classical streamline-based tractography techniques [11] as illustrated in Figure 1(c). From a seed point *P*_0_ = (*x*_0_,*y*_0_, *z*_0_), a new streamline is created with the initial seed 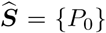. Next, the model is given the DWI data at the previous streamline coordinate *P_i_* to obtain a predicted direction 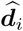. A next point 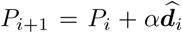 is then computed, where α is a chosen step size, as in standard streamline-based tractography algorithms. Points are generated to iteratively produce a streamline until a desired criterion is met (e.g. too high curvature, exiting WM mask). The whole process is repeated as many times as required to produce a full tractogram.

## 3 Related Work

The work of Neher *et al.* [13] hypothesizes that tractography can be improved by considering local neighborhood features, adding a directional prior to promote straight fibers, and using a fiber deflection protocol to help the model recover from mistakes. More precisely, their model makes predictions based on a voting mechanism, using local direction proposals from multiple sample positions in the vicinity of the current location. Each direction proposal is obtained by a classification over 100 possible directions (weighted using the previous direction), along with a streamline termination probability. If fiber termination is the more likely option, a deflection is attempted by rotating the sample point 180° around the previous direction and classifying a second time.

In our current approach, the problem is framed as a regression task over normalized directions instead of a vote over discretized directions. This means that to produce a prediction, fewer computations are needed at the output, compared to computing and voting over many proposals. Regression also allows the model to output more precise directions and thus be more suitable at exploiting smaller variations in direction. In addition, if straight fibers are supported by the data, a directional prior should not be necessary and a deep learning model should be able to learn the right structure, which is why our model does not include such a prior.

While Neher *et al.* consider the neighborhood of the current position, they do not consider the full evolution of the streamline up to each point. Our hypothesis is that there are high-order dependencies between the next direction in the streamline and all previous directions. Consequently, our recurrent approaches have a natural mechanism for integrating past information along the streamline to predict a next direction. These two approaches are not exclusive however, and would probably benefit from each other.

Finally, in a deep learning context, learning a stopping criterion along with the direction to follow is more complex. It would require careful engineering and balancing of the loss function in order for one not to overcome the other during training, especially using a recurrent approach. This is beyond the scope of this paper and left for future work.

## 4 Experiments

We quantify the performance of our methods on the 2015 ISMRM challenge dataset [10] and evaluate using the Tractometer connectivity metrics [4]. In doing so, we can compare ourselves to the 96 original challenge submissions [1]. We then qualitatively evaluate our method when tracking on *in vivo* data.

*Tracking parameters:* Tracking was done using ½ voxel step size (1.0 mm for ISMRM challenge, 0.625 mm for HCP). Seeding was done using 1 seed per voxel in the WM mask, and tracking was done using a dilated WM mask. All streamlines leaving the dilated mask were automatically terminated and streamlines shorter than 20 mm or longer than 200 mm were discarded. Streamlines with a half-cone curvature higher than 20° were also discarded.

### 4.1 ISMRM2015 Challenge

For the first experiment, we choose to reproduce the training environment of Neher *et al.* [13], by using training data generated specifically for the subject of interest. They determined the optimal method for generating their training data to be a constrained spherical deconvolution (CSD) deterministic streamline tractography (DET) of MRtrix [14]. Staying coherent with their approach, a tractogram was generated using CSD-DET on a denoised and distortion-corrected version of the ISMRM2015 challenge dataset. The resulting 92K streamlines were then split into training and validation sets (using splits of 90% and 10%).

We trained models with one to four layers, varying the layer size between 500 and 1000, and used the Adam optimizer with early stopping. Full code is available online^5^. For each type of model, the one with the best validation error was chosen for tracking & tractometer evaluation.

We report in Table 1 the valid connections ratio and the associated number of valid bundles (true positives), the invalid connections ratio and the associated number of invalid bundles (false negatives), the volumetric bundle overlap and overreach in percentages. Drawing conclusions from only one of these last two metrics can be misleading (e.g. a model can have both the best overlap and the worst overreach). These metrics are related to precision and recall measures, and are combined into the F_1_-measure. Note that the “_PD” model suffix indicates when the previous direction was given as input to the model. We report as baselines the ISMRM mean results and submission 6_1[1], which is a CSD-DET based method comparable to what was used to generate our training data.

We see that the local model (FFNN) is already competitive with the mean ISMRM challenge scores. Its ability to estimate the main diffusion axis and tuning its predictions according to the streamlines seen during training allows it to improve all mean scores except number of invalid connections (+3.9 VC, +7.6 IC, −10 NC, +1.6 VB, −214 IB, +12.2 OL, −1.6 OR, +11.6 F1). Surprisingly, adding the previous direction as input worsened the model's performance. We think that the optimization process allowed the model to achieve a good performance by generally simply copying the previous direction given as input. Indeed, we looked at the generated streamlines, and while they do a good job of covering the brain (the FFNN_PD model recovered 22 bundles out of 25), they are mostly straight and miss important connections.

**Table 1.**
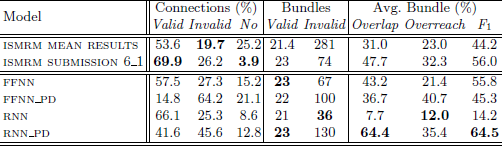
Quantitative evaluation on the ISMRM 2015 Tractography Challenge.

**Fig. 2.**
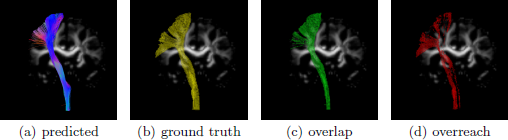
a) Left CST generated by the RNN model using the ISMRM2015 challenge data. b) Ground truth mask as defined by the ISMRM2015 challenge. c) Overlapping and d) overreaching voxels of the generated bundle with respect to the ground truth mask.

Going from the local model to the recurrent model (RNN) provided different insights. Without the previous direction as input, the model generated more relative valid connections, but overall very few streamlines (as seen in the overlap metric, 7.7%). With the previous direction however, the model achieved very good coverage of the challenge bundles (64.4% overlap), while dropping a bit below 50% VC. It achieved the best F1 score over all our models. **In comparison, no submission in the ISMRM2015 challenge achieved an overlap higher than 50% while keeping overreach under 50%** [1,10]. Figure 2 shows how the left CST is reconstructed with high coverage and low overreach. We believe that the recurrent model, being able to accumulate “memories” about the past of the streamline, is able to extract information of the previous direction without committing the same mistakes as the local model. This ability to “memorize” is what makes this model stand apart from classic methods.

**Fig. 3.**
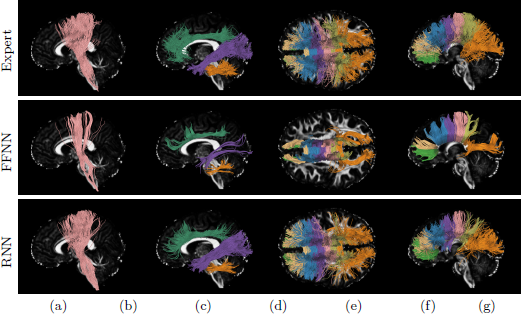
Expert, FFNN and RNN bundles obtained in experiment 4.2. Colored bundles (in order) : [CST], [Cingulum, ILF, MCP], [Rostrum, Genu, Rostral Body, Anterior Midbody, Posterior Midbody, Isthmus, Splenium].

### 4.2 In *vivo* tracking

Using the models trained in the first experiment (section 4.1), we tracked on an unseen brain (HCP subject #100307). As a gold standard we used a virtual ROI-based dissection made by an expert neuroanatomist [12,2]. Streamlines used for the dissection were generated using Particle Filtering Tractography [6] using default parameters and based on a spherical harmonics 8 multi-shell constrained spherical deconvolution reconstruction [5,8]. The resulting bundles are shown in Figure 3. Visual evaluation shows results that are in line with the first experiment. The local model does a good job of recovering the bundles, but has poor coverage. The recurrent model is much more similar to the expert segmentation in most of the recovered bundles. We suspect that the RNN would gain even more by training on much larger datasets with multiple subjects.

## 5 Conclusion

We propose the first deep learning alternatives to traditional local modeling approaches to tractography based on raw DWI. Our FFNN model provides the first performance baseline for local deep models. We also present a novel approach where the past of the streamline is considered by a recurrent model in order to make better predictions. Compared to the other ISMRM2015 submissions, this proved to be the only technique able to recover more than 50% of spatial coverage while producing overreaching false connections below 50%. We show that deep learning models can generalize to new DWI unseen at training time. These novel results show that deep learning is a promising approach to tractography.

While we believe that deep learning will be able to discover new pathways by learning the global streamline structure, we still do not have enough accurate data to explore this area of research. In future works, as data become available, we plan on training on incomplete datasets (i.e. removing one or more bundles) in order to see the reconstruction and discovery capabilities of our models. Furthermore, we will explore how modifying the output of the RNN (e.g. predicting the parameters of a distribution) can improve the power of the model.

## Footnotes

5 https://github.com/ppoulin91/learn2track/tree/miccai2017_submission

